# From Prompt to Provenance: BloClaw, a Capability-Gated AI4S Workstation for Auditable Computational Biology

**DOI:** 10.64898/2026.08.26.747436

**Authors:** Yao Qin, Jinhua Pang, Xiaoming Zhang

## Abstract

Scientific agents can produce plausible answers while remaining unable to establish whether the computation behind an answer is executable, recoverable, or reproducible. We present BloClaw, an AI4S workstation built around a simple principle: a scientific agent should know what it can do, show how it did it, and state what remains unvalidated. Each capability declares an execution state, input constraints, dependencies, expected outputs, and scientific limitations. Natural-language requests are translated into structured tasks, validated against this registry, executed through scientific tools, and recorded in a provenance-aware Living Lab Notebook. The system is designed to detect invalid inputs, failed tool calls, missing dependencies, and remote timeouts, and to route them to repair, retry, or escalation. The implemented and tested scope comprises RDKit-based molecular property and rule screening, protein structure analysis, docking-pose inspection, 3D visualization, and structured reporting. We demonstrate the workflow on a PubChem-retrieved osimertinib structure and a supplied 6LU7 docking artifact: the former yields deterministic descriptors (molecular weight 499.619 Da, cLogP 4.5098, TPSA 87.55 A^2^), while the latter contains 2,387 protein ATOM records, 309 residues, and nine pose records. These examples are workflow demonstrations, not efficacy or affinity studies. Beyond retrospective prediction, the manuscript specifies a prior-minimized constructive mode in which a desired function is compiled into explicit physical, chemical, and systems constraints, candidate mechanisms are simulated, and observations are reintroduced for calibration and falsification; this is a proposed extension rather than a result of the present case studies. We describe an evaluation protocol that compares BloClaw with a standard single-agent workflow and fixed-script execution using task completion, scientific correctness, recovery success, provenance completeness, reproducibility, human review time, latency, and cost. This manuscript reports the system design, verified capability boundary, deterministic software artifacts, and a reproducible evaluation protocol; it does not claim benchmark improvements before those experiments are run. BloClaw is an execution and accountability layer for AI-assisted research, complementing expert review and experimental validation rather than replacing them.

## 1 Introduction

Large language models (LLMs) have made natural-language interaction with computational tools accessible to a broader range of researchers. Recent systems have demonstrated LLM-guided chemistry-tool use and partially autonomous laboratory planning, while also illustrating the need for explicit tool and execution controls [1, 2]. In computational biology, an LLM can formulate analysis plans, select software, interpret intermediate results, and prepare reports. Fluent language generation, however, is not equivalent to reliable scientific execution.

The central question is not only whether an agent can generate a plausible answer, but whether the underlying scientific action is executable, recoverable, and auditable. A useful system must verify that a capability is actually available, validate its inputs, preserve the parameters and software versions used, recover from tool failures, and distinguish measured evidence from model-generated interpretation. Scientific-agent workflows can therefore fail in four recurring ways: capability over-claiming, invalid or incomplete tool execution, missing provenance, and overinterpretation beyond an appropriate applicability domain.

We present BloClaw, an AI4S workstation designed to make the capability boundary an executable part of the workflow rather than a descriptive claim in the user interface. The system accepts natural-language research requests, translates them into structured tasks, validates capabilities and inputs, orchestrates scientific tools, records evidence and uncertainty, and produces exportable research artifacts. The capability matrix in the current repository reports 3 functions as `available`, 36 as `migrating`, and 29 as `planned`. This paper treats only the available functions as validated scientific scope.

The research question is:

Can capability-gated execution and structured provenance reduce silent failures and human verification burden in computational biology tasks, compared with a standard single-agent workflow and fixed-script execution?

The contributions are:

- a capability-state registry that turns executable, migrating, and planned functions into explicit task constraints;
- a recoverable task-execution lifecycle for computational biology workflows;
- a provenance-aware Living Lab Notebook linking hypotheses, inputs, tools, parameters, outputs, and evidence;
- a scope-controlled set of molecular, structural, docking-pose, and reporting artifacts that can be inspected and rerun;
- a benchmark and ablation protocol for testing scientific correctness, recovery, provenance completeness, reproducibility, and human review burden;
- a prior ledger and constructive-mode specification that separates axioms, measurements, executable models, and human objectives, with tests for prior leakage and out-of-distribution mechanism generation.

## 2 Related Work

### 2.1 LLM-based scientific agents

LLM-based agents have been used to plan analyses, call external tools, write code, and interpret scientific results [1, 2]. Their flexibility is useful for exploratory work, but reliability depends on tool interfaces, validation, and execution control. BloClaw focuses on the execution and accountability layer surrounding an LLM rather than on a new foundation model.

### 2.2 Scientific workflow systems

Workflow managers provide deterministic pipelines, dependency handling, and execution logs. Representative systems include Snakemake, Nextflow, and Galaxy [3, 4, 5]; notebook, container, and provenance practices further support reproducible execution [6, 7, 8, 9, 10, 11]. They are effective when a workflow is known in advance, but generally require researchers to specify it explicitly. Blo-Claw adds natural-language task interpretation while retaining explicit execution and validation boundaries.

### 2.3 Provenance and reproducible research

Scientific provenance systems record relationships among data, software, parameters, and outputs. The FAIR principles provide a widely used framework for making research objects findable, accessible, interoperable, and reusable [12]. Reproducible computational research also requires explicit control of inputs, software, parameters, and execution environments [13, 7, 6, 9, 8]. Formal provenance models and workflow surveys provide complementary representations for linking activities, entities, and agents [10, 11]. BloClaw combines these ideas with an interactive AI4S task interface and a structured research notebook.

### 2.4 Research gap

Many existing systems emphasize either conversational interaction or deterministic workflow execution. The gap addressed here is the combination of natural-language interaction, capability-state gating, recoverable tool execution, and provenance-aware scientific delivery in one operational workstation. The distinction is architectural and operational; it is not a claim that BloClaw replaces established workflow systems.

### 2.5 From retrospective prediction to constructive AI4S

Most scientific AI systems are evaluated as predictors over a historically populated data manifold: observations are retrieved, a representation is learned, and a likely output is interpolated or extrapolated from related examples. This paradigm is useful for measurement-rich tasks, but it is poorly matched to a living organism whose response depends on transport, feedback, adaptation, tissue context, and time. A molecule can satisfy a target-level assay while failing to move the organism along a safe, therapeutic trajectory.

BloClaw therefore specifies a complementary *constraint-first constructive mode*. At task initiation, an epistemic reset separates four sources of authority: explicit axioms, measurements, executable models, and human objectives. Unverified retrieval context is withheld from candidate generation; the language model is restricted to parsing and orchestration; and a declared mechanism kernel composes candidates from physical, chemical, and systems-level constraints. This task-level prior isolation does not erase pretrained weights or claim a literal blank slate; it makes the allowed sources of candidate novelty explicit and testable. Historical measurements are then reintroduced to calibrate uncertain parameters, discriminate competing mechanisms, and falsify predictions. This is not data-free science. It is data-independent hypothesis generation followed by data-dependent validation.

The constructive task can be written as *T* = (*G, A, C, O*), where *G* is a desired function or phenotype, *A* is a set of explicit mechanistic axioms, *C* is an engineering and biological constraint set, and *O* is optional observational data. A candidate mechanism *x* is generated from a declared space *X* (*A, G*) and evaluated as

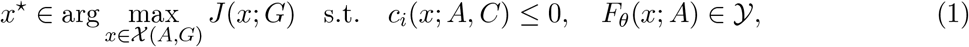

where *F*_*θ*_ is an executable simulator and law parameters are kept separate from empirically calibrated parameters. For life-design tasks, the minimum specification includes boundary integrity, energy transduction, catalytic closure, information storage and replication, error control, homeostasis, perturbation recovery, and evolvability. Creator’s-eye design is used only as a metaphor for starting from these invariants; it is not a claim of omniscience or autonomous life creation.

### 2.6 An operational ontology of life-like systems

BloClaw does not claim a universal philosophical definition of life. For design and simulation, it uses an operational criterion for an autonomous protocell-like system in an environment over a finite horizon. The system is treated neither as source code alone nor as a disembodied information projection: code describes possible transitions, while information has biological meaning only when a physical state changes a future trajectory. The vector is reported component-wise rather than collapsed into an arbitrary score. For a single bounded candidate, *B, E, M* , *I*, and *R* are hard design invariants; *V* is evaluated at the lineage or population level, or marked not applicable for a deliberately non-evolving construct:

- *B* (boundary persistence): compartments or interfaces remain sufficiently intact to regulate exchange;
- *E* (energy transduction): external free energy is converted into maintained gradients and useful work with entropy export;
- *M* (self-maintenance): catalytic and repair processes replenish the components needed to keep the system running;
- *I* (information copying): a heritable information state is stored and copied with a measurable error rate;
- *R* (perturbation recovery): the system returns to a declared viability region after specified disturbances;
- *V* (evolvability): constrained variation can change function without destroying the preceding conditions.

The candidate is called *life-like* only when the applicable components exceed preregistered thresholds under a finite resource budget. Boundary cases such as viruses, dormant forms, and obligate symbionts are reported with their dependency assumptions rather than forced into a binary label. This is an engineering test for a virtual or synthetic system, not a claim that these design invariants settle the biology or philosophy of life. A useful state-space view is the viability kernel: the set of initial states from which some admissible control policy keeps the mechanism inside the declared constraint set while achieving the task within the time window. Its estimated volume, recovery time, energy cost, and copying fidelity become measurable design objectives.

## 3 System Design

### 3.1 Design goals

BloClaw was designed around six goals:

1. capability honesty: do not expose an unavailable computation as completed;
2. scientific input validation: check files, formats, dependencies, and parameter constraints before execution;
3. recoverable execution: detect, repair, retry, or escalate failures;
4. provenance completeness: preserve the information required to reconstruct a result;
5. human-verifiable delivery: expose intermediate outputs, limitations, and uncertainty to the researcher;
6. epistemic separation: distinguish axioms, measurements, executable models, human objectives, and speculative proposals.

### 3.2 Four-layer architecture

The system is organized into four cooperating layers (Figure 1).

**Figure 1:**
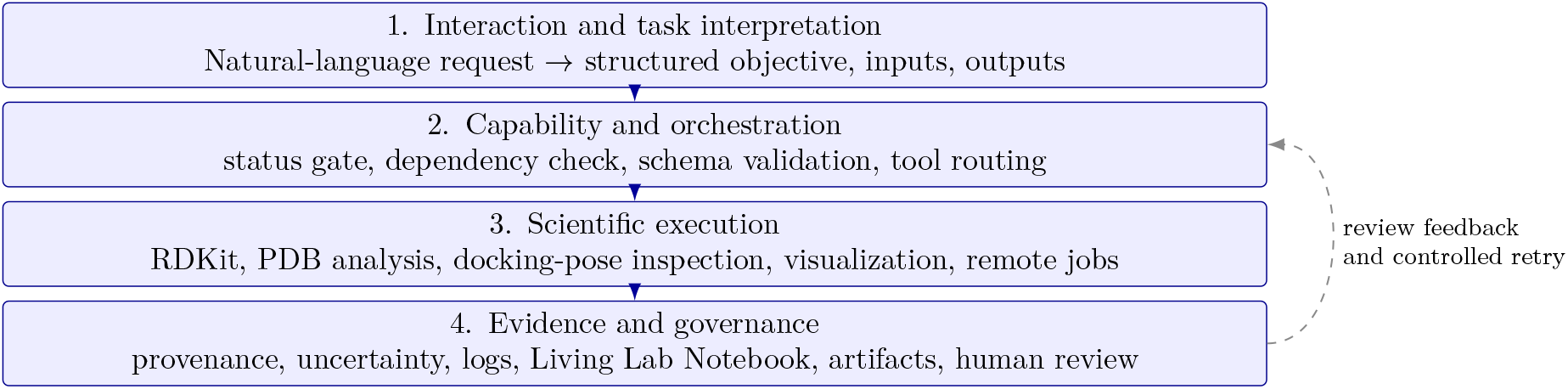
BloClaw four-layer architecture. Natural-language interaction is constrained by a capability registry before scientific tools are executed; outputs enter the evidence and governance layer for provenance, uncertainty, review, and export.

#### Interaction and task interpretation

Natural-language requests are converted into structured objectives, required inputs, expected outputs, and review criteria. The structured task is stored with the run rather than inferred retrospectively from a chat transcript.

#### Capability and orchestration

The capability registry records status, dependencies, accepted input formats, expected outputs, failure conditions, and scientific limitations. The orchestrator selects tools and routes subtasks only after checking these constraints.

#### Scientific execution

This layer executes validated molecular, structural, visualization, and remote-compute operations. Each operation returns structured outputs and execution metadata.

#### Evidence and governance

This layer records provenance, uncertainty, task logs, notebook entries, project boundaries, and exportable artifacts. It communicates when a capability is unavailable or when an output requires human review.

#### 3.3 Capability-state gating

Every capability is assigned one of three states:

- `available`: implemented, dependency-checked, and tested for the declared task scope;
- `migrating`: partially implemented or under integration, and not suitable for strong scientific claims;
- `planned`: not currently executable.

The state is an execution constraint, not merely a user-interface label. A request that exceeds the declared scope is rejected, downgraded to an explanatory response, or escalated for human review (Figure 2).

**Figure 2:**
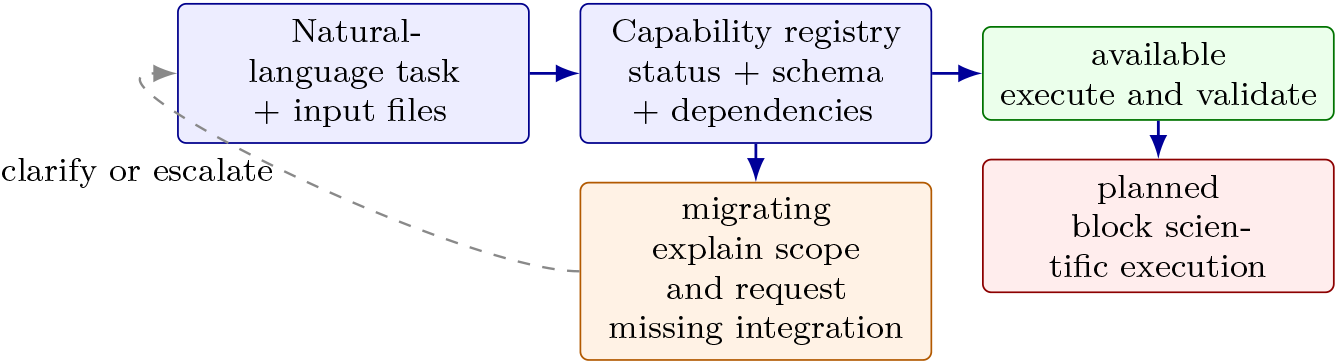
Capability-state gating. The same request is routed differently depending on registry state; only an available capability reaches scientific execution.

**Figure 3:**
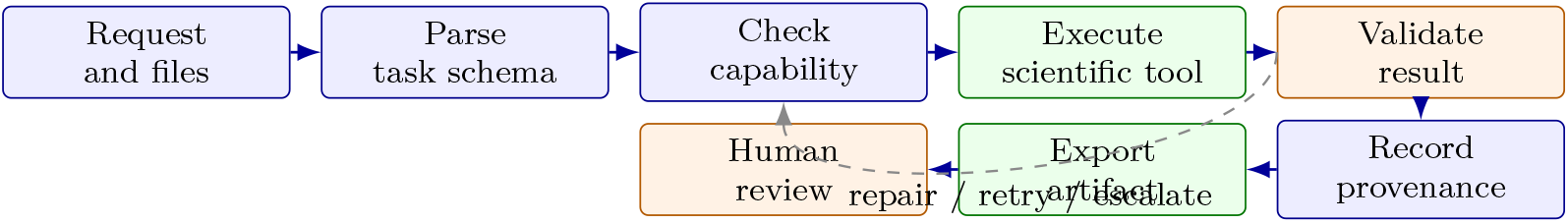
Task-execution lifecycle. Failures detected after result validation return to the capability check for controlled repair, retry, or escalation.

### 3.54 Task-execution lifecycle

The lifecycle consists of request parsing, input extraction, capability validation, tool and parameter selection, execution, error detection, recovery or escalation, result collection, provenance recording, and human review. Each stage produces a machine-readable event that can be inspected after completion.

### 3.5 Failure recovery

The recovery subsystem handles invalid or missing files, unsupported formats, missing dependencies, incorrect parameters, remote timeouts, partial outputs, and visualization-loading failures. A recovery event records the original error, corrective action, re-execution status, and whether the final output passed validation. Recovery is not allowed to silently substitute demo data for a failed scientific step.

### 3.6 Provenance and Living Lab Notebook

The Living Lab Notebook links a hypothesis to input files, tools, versions, parameters, intermediate outputs, final results, uncertainty notes, human decisions, and evidence links. It is a structured scientific record rather than a conversation archive (Figure 4).

**Figure 4:**
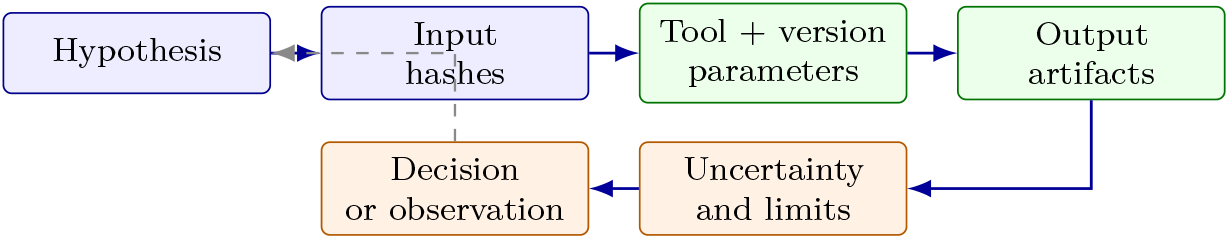
Provenance graph represented by the Living Lab Notebook. Evidence flows from a hypothesis through hashed inputs, named tools and parameters, output artifacts, uncertainty, and a recorded decision or observation.

**Figure 5:**
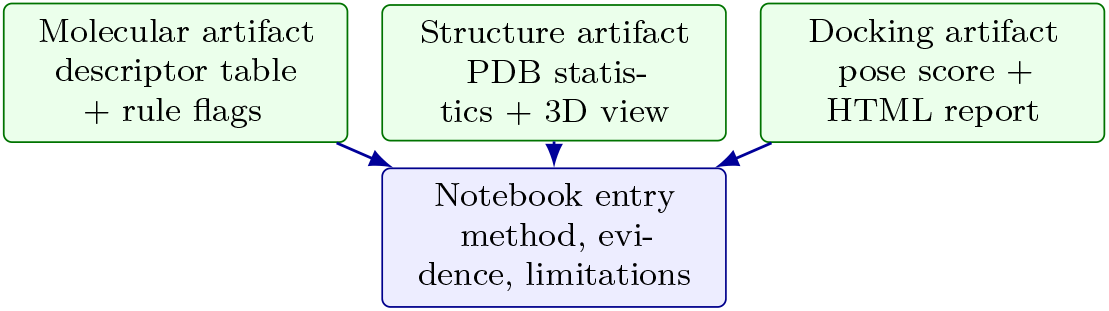
Representative scientific deliverables produced by the validated scope: a molecular descriptor table, a PDB structure report with 3D view, a docking-pose report, and a linked Notebook entry.

## 4 Implemented Scientific Scope

### 4.1 Molecular property and rule screening

The validated molecular workflow ingests SMILES or molecular files, computes RDKit descriptors, applies rule-based filters, and generates tabular outputs with source and parameter metadata. The workflow is explicitly a physicochemical and rule-screening function; it is not complete ADMET prediction. The rule checks are interpreted in the context of established drug-likeness and solubility heuristics [14, 15, 16]. For the case study below, the canonical SMILES for osimertinib was retrieved from PubChem CID 71496458 [17], then processed by the same deterministic helper used by the workstation.

### 4.2 Protein structure analysis

The structural workflow parses Protein Data Bank (PDB) files, reports chains, residues, atoms, ligands, and structural anomalies, and exports a visual inspection artifact. The PDB is a primary archive for experimentally determined three-dimensional biological structures [18]. Modern predictors and structure databases such as AlphaFold, ESMFold, RoseTTAFold, RFdiffusion, and AlphaFold DB illustrate adjacent capabilities but are not claimed as implemented here [19, 20, 21, 22, 23]. The declared scope is structure analysis and quality inspection, not protein-folding prediction.

### 4.3 Docking-pose inspection and visualization

The docking workflow reads existing pose files, reports scores and pose identifiers, renders protein– ligand views using WebGL-based molecular visualization [24], and exports an HTML report. Molecular simulation and scientific-computing libraries provide relevant execution context for reproducible numerical workflows [25, 26, 27, 28, 29, 30]. AutoDock Vina is cited only when its output format or score is part of the input artifact [31]; pose inspection does not establish binding affinity, biological activity, or therapeutic efficacy.

## 5 Empirical Case Studies

### 5.1 Osimertinib physicochemical triage

The PubChem canonical SMILES for osimertinib (formula C_28_H_33_N_7_O_2_) was processed with RD-Kit in the pinned BloClaw helper. The output was molecular weight 499.619 Da, cLogP 4.5098, topological polar surface area 87.55 A^2^, two hydrogen-bond donors, seven acceptors, ten rotatable bonds, 37 heavy atoms, and fraction sp^3^ of 0.25. The Lipinski-style and Veber-style checks implemented by the helper returned no flags (Figure 6). This is a deterministic property report and rule triage; it does not estimate CYP, hERG, Ames, permeability, clearance, or clinical toxicity.

**Figure 6:**
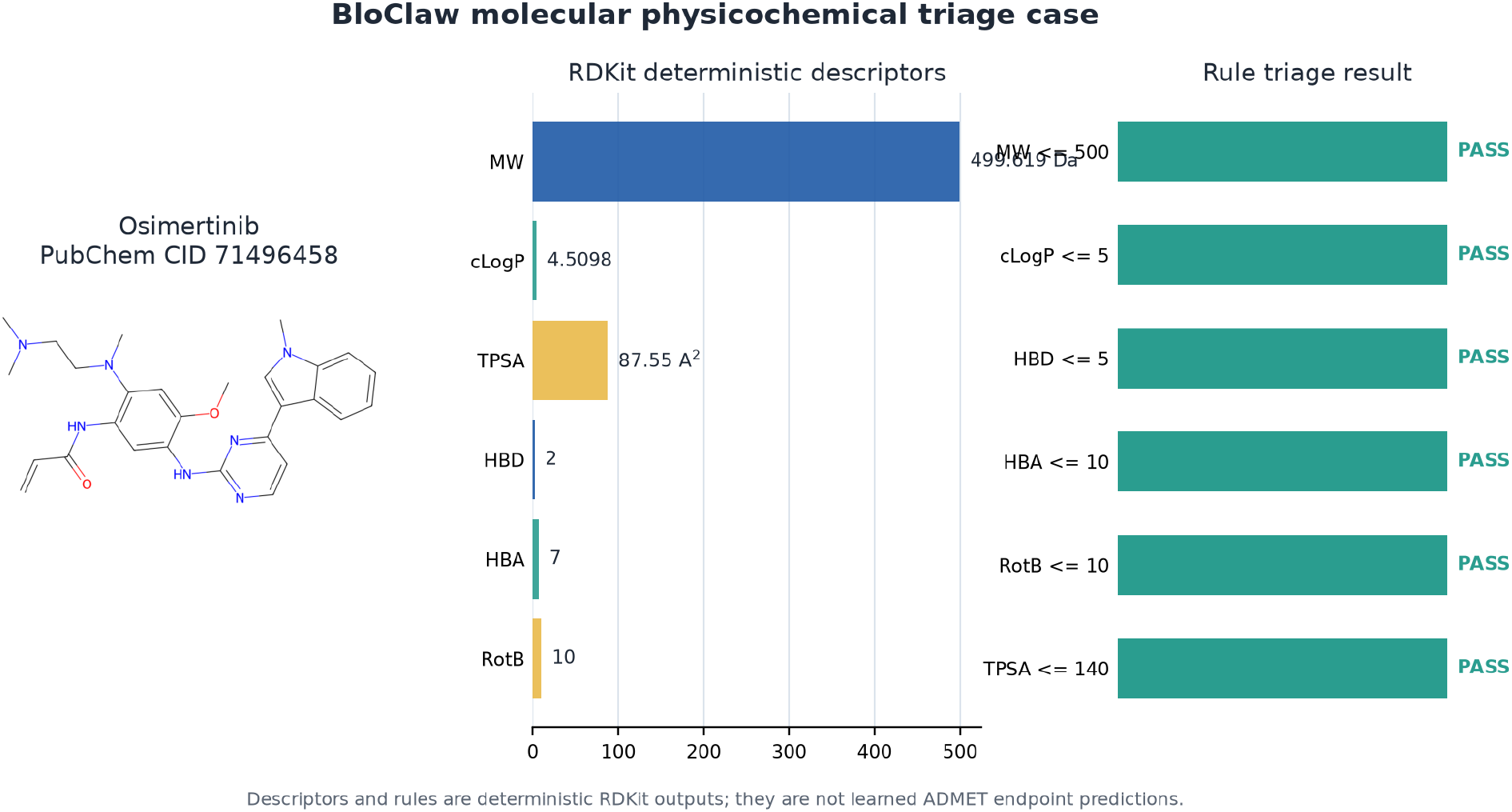
Osimertinib molecular physicochemical triage. The structure is rendered from the Pub-Chem CID 71496458 canonical SMILES and the descriptor and rule panels are generated from the BloClaw RDKit helper. The pass labels refer only to the implemented threshold checks and should not be interpreted as ADMET or clinical predictions.

### 5.2 6LU7 docking-pose inspection

The supplied BloClaw HTML artifact contains an embedded protein structure and nine ligand pose records. Parsing the embedded data produced 2,387 protein ATOM records, 29 HETATM records, 309 residue identifiers across chains A and C, and 39 atoms in each pose. The viewer reports score fields of 8.01, 7.87, 7.80, 7.78, 7.63, 7.58, 7.56, 7.56, and 7.26 kcal/mol for poses 1–9, respectively (Figure 7). These are the values displayed by the supplied artifact; because the source does not include an independently rerun scoring calculation or a signed Vina log, we report them as displayed score fields rather than as measured binding free energies. The protein and ligand coordinates are shown for inspection only, and no affinity, efficacy, or mechanistic claim is made.

**Figure 7:**
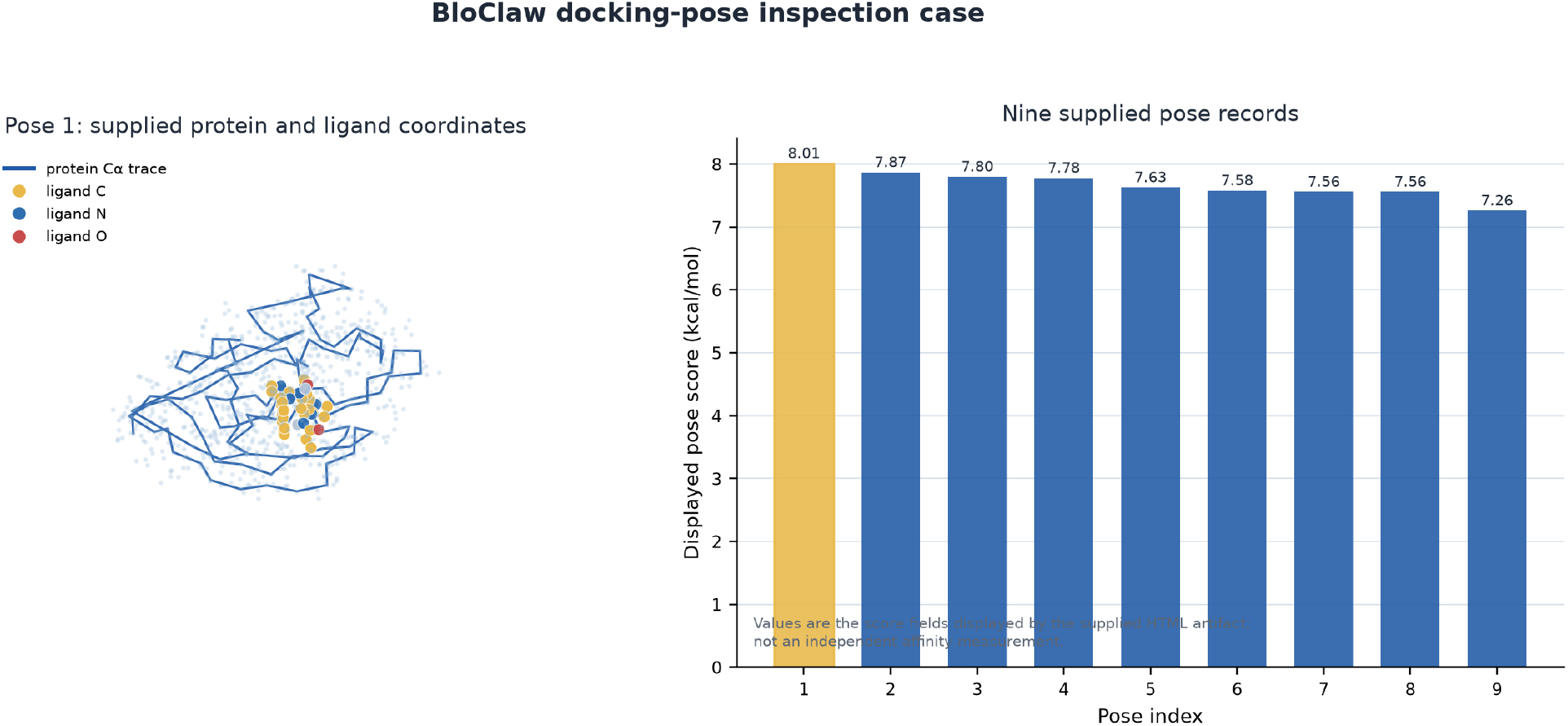
Docking-pose inspection case based on the supplied BloClaw HTML artifact. Left: protein C*α* trace and Pose 1 ligand coordinates after centering on the ligand; the pale points show nearby non-hydrogen protein atoms. Right: nine score fields displayed by the artifact. The figure documents file parsing and visualization, not an independent docking calculation or biological validation.

**Figure 8:**
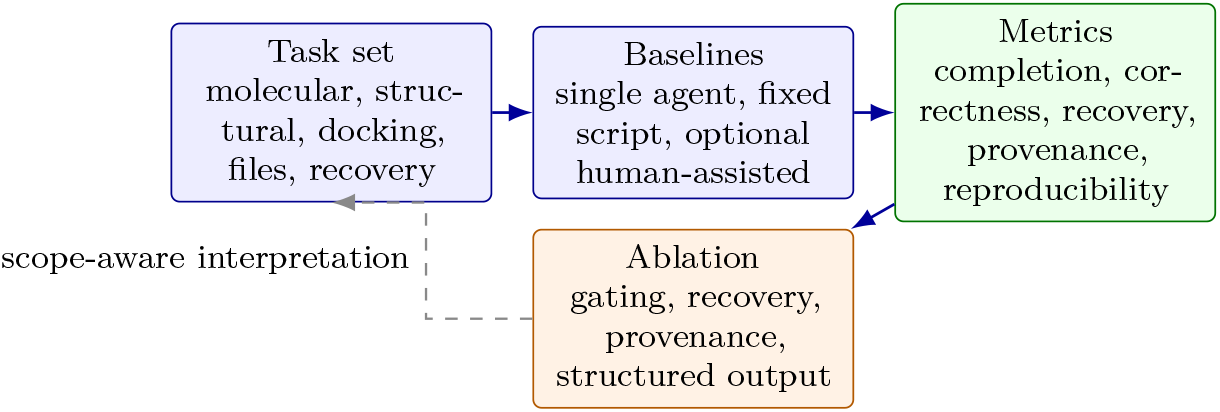
Evaluation protocol for the next empirical stage. The present manuscript makes no quantitative superiority claim; the figure defines the task, baseline, metric, and ablation structure required to test one.

### 5.3 Artifact-level validation

The two cases exercise different validated paths: deterministic molecular descriptors and parsing/inspection of supplied structural artifacts. Their outputs are stored with the source identifiers, method description, and limitations. The docking source artifact has SHA-256 digest `0074968680b18ec4719a7ad1ac6e8eb453f3a9d7ee3f7f6ec1157fcc2e3e3ca4;` the extracted case-data manifest and figure-generation scripts are included with this manuscript package. These checks establish that the figures can be regenerated from the named inputs; they do not establish biological performance.

### 5.4 Remote execution and isolation

Remote tasks are assigned identifiers and tracked through submission, running, completion, failure, timeout, recovery, and collection states. Project-level sandboxing limits file access and separates input data, intermediate files, and exported results. Remote jobs can be restart-recoverable, but full scientific provenance for every remote dependency remains a stated implementation boundary and is not claimed as complete here.

## 6 Software Validation and Evaluation Protocol

### 6.1 Verified capability declaration

The repository capability matrix reports three available capabilities and separates them from migrating and planned functions. Table 1 records the manuscript treatment.

**Table 1:** Capability declaration used in this manuscript.

| Capability | State | Evidence and claim boundary |
| --- | --- | --- |
| Molecular property and rule screening | available | RDKit descriptors and Lipinski/Veber-style rule checks; not complete ADMET |
| Protein structure analysis | available | User-provided PDB parsing and inspection; no folding prediction |
| Docking-pose inspection and visualization | available | Existing pose review and 3D report; no de novo docking claim |
| Complete ADMET prediction | planned | Excluded from validated results |
| De novo molecular docking | planned | Excluded unless a declared engine is independently benchmarked |
| Protein folding and virtual-cell perturbation | planned | Architecture extension only |
| Antibody, radiopharmaceutical, peptide models | migrating/<br>planned | Future work unless independently evaluated |
| Autonomous robotic execution | planned | Outside the scope of this manuscript |

### 6.2 Deterministic software checks

The current validation package should be understood as an artifact-level software check, not as a biological efficacy study. For molecular screening, deterministic descriptor values are checked against a pinned RDKit environment. For structure analysis, the parser output is checked against atom, residue, chain, and ligand counts derived from the same PDB input. For docking-pose inspection, pose identifiers, score fields, input-file hashes, and exported HTML assets are checked for integrity. For every artifact, the corresponding Notebook entry records the method, inputs, parameters, limitations, and output path.

### 6.3 Benchmark task set

The next empirical stage should contain 20–50 tasks across empirical retrieval and screening, PDB structure analysis, docking-pose inspection, mechanistic derivation, constructive candidate generation, active-experiment selection, failure injection, and repeated execution. Every task must have an input package, expected output, acceptance criteria, and a reference computation or expert-verified answer. Constructive tasks should use held-out or synthetic worlds with randomized names and disabled retrieval so that language-model memory cannot define the candidate space. The present case studies are not used as a benchmark score or baseline comparison.

### 6.4 Baselines and metrics

The primary comparison should include a standard single-agent or retrieval-augmented workflow, a fixed constrained sampler, and a fixed-script workflow using the same scientific tools and inputs. A human-assisted workflow may be added to quantify review burden. In constructive tasks, metrics must additionally include constraint-satisfaction rate, mechanistic consistency, candidate diversity and novelty under a declared audit, out-of-distribution viability, falsification efficiency, calibration, prior leakage, assumption traceability, and information gain per experiment. All tasks must also report task completion, scientific correctness, tool-call error rate, recovery success, provenance completeness, reproducibility, human review time, latency, and execution cost. Define every metric before testing and report confidence intervals for repeated runs.

### 6.5 Ablation and reproducibility protocol

The ablation study removes capability gating, failure recovery, provenance recording, and structured-output constraints one at a time. For constructive tasks, additional ablations remove retrieval suppression, mechanistic constraints, the prior ledger, and active feedback; a language model restricted to parsing and planning is compared with a model allowed to propose mechanisms directly. The evaluation must record model and provider versions, generation settings, software versions, dependency environment, hardware or remote runtime, inputs, prompts, parameters, random seeds, logs, and output files. This follows established recommendations for FAIR and reproducible computational research [12, 13].

## 7 Case Studies and Scientific Boundaries

### 7.1 Molecular screening case

The osimertinib case above reports molecular weight, lipophilicity, hydrogen-bond donors and acceptors, rule-based alerts, a screening decision, and provenance metadata. The result is a transparent physicochemical triage artifact and should not be described as a prediction of clinical exposure or toxicity.

### 7.2 Protein structure case

The 6LU7 artifact reports file identity, chains, residues, atoms, hetero records, and a linked three-dimensional view. It is an inspection of an input structure, not a prediction of an unobserved fold.

### 7.3 Docking-pose case

The existing docking artifact reports pose ranking, score fields, pose metadata, protein–ligand visualization, and a reproducible HTML export. The 6LU7 + osimertinib example is a software workflow demonstration only; it is not evidence of efficacy, affinity, or a new biological mechanism.

### 7.4 Failure-recovery case

A controlled failure test is specified for the next evaluation stage. It should show an invalid input, detected problem, corrective action, re-execution, and final validated output. If the problem cannot be repaired within the declared scope, the correct output is an explicit blocked or escalated step; no failure-recovery success rate is claimed in the present paper.

## 8 Proposed Constructive Extension

The validated cases in this manuscript demonstrate artifact-level execution and accountability. The next scientific extension is to make the same execution contract work for constructive, mechanism-first design. The unit of design is then not an isolated molecule or a single endpoint score, but an intervention trajectory in an embodied, multiscale living system. A candidate must preserve a declared viability region while moving the system toward a specified function under uncertainty, resource limits, and perturbations.

### 8.1 Constraint-first life-design loop

The proposed loop begins with a target function, phenotype, environment, and resource budget. A versioned axiom compiler translates these requirements into conservation, thermodynamic, chemical, geometric, and boundary constraints. A typed mechanism grammar then composes reaction, interaction, regulatory, and compartment modules. Candidates are statically checked, simulated at the appropriate scales, and stress-tested under counterfactual environments. An active experiment planner selects the smallest set of observations that can discriminate competing mechanisms. Measurements update uncertain parameters or reject a mechanism; they do not silently rewrite the original hypothesis. Figure 9 summarizes this proposed extension.

**Figure 9:**
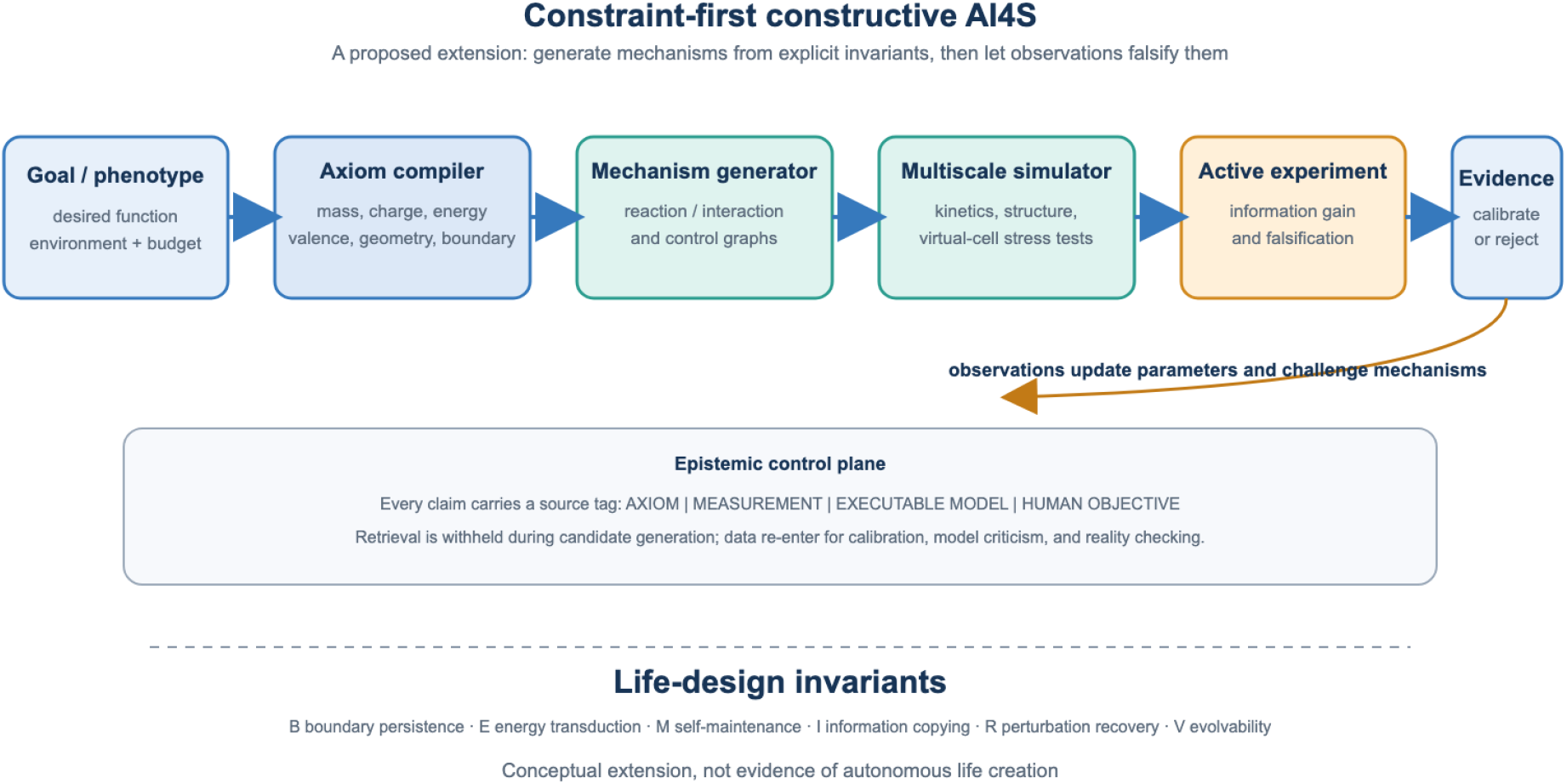
Constraint-first constructive AI4S loop. A desired function is compiled into explicit invariants and constraints before a mechanism is generated. Simulation and active experiments provide calibration and falsification evidence. The loop is a proposed extension and is not evidence of autonomous life creation.

### 8.2 Candidate design targets

For antibody therapeutics, constructive generation would start from epitope geometry, kinetic objectives, folding stability, aggregation limits, immunogenicity constraints, expression feasibility, and tissue exposure, then generate and simulate sequence–structure mechanisms before selecting discriminating assays. This is materially different from ranking sequences only by similarity to known antibodies; it is also outside the validated capability scope of the present manuscript.

For radiopharmaceuticals, the specification would jointly constrain radionuclide half-life and decay energy, chelator coordination, linker stability, target binding, biodistribution, clearance, and absorbed-dose limits. The output would be a molecule–schedule–tissue trajectory with uncertainty, rather than a single affinity score. No radiopharmaceutical prediction is claimed here.

For virtual-cell design, the primitive objects would be compartments, metabolites, catalysts, transporters, information states, and feedback rules. A candidate network would be required to satisfy mass and energy balance, non-negative concentrations, self-maintenance, perturbation recovery, and an explicitly defined information or replication task in an unseen environment. A reduced simulator can test these properties, but it must not be presented as a whole-cell or patient-specific predictor without independent validation.

### 8.3 A causal failure model for AIDD translation

The problem is not that every AIDD system fails, but that a high score on a historical assay is not the same object as a therapeutic effect in a changing organism. A recurrent failure chain starts with selection and measurement bias in historical data, proceeds through shortcut features and proxy optimization, and then encounters scaffold, assay, species, tissue, time, or synthesis shifts at deployment. Static endpoint models can additionally omit transport, active metabolites, feedback, adaptation, toxicity, and resource limits. Selective wet-lab follow-up then feeds only a subset of outcomes back into the training set, creating data drift and confirmation bias. A language model can add a second failure channel by leaking familiar names or pathways into candidate generation when the evidence is missing. BloClaw’s proposed intervention points are explicit hard constraints before optimization, mechanism and trajectory simulation, counterfactual perturbation tests, active experiment selection, and evidence-level auditing. This is a failure hypothesis to be tested, not a claim that correlation-based AIDD has no value.

### 8.4 Falsifiable predictions of the constructive mode

The constructive mode makes prospective predictions rather than asserting that a new theory is already proven. Under matched compute, candidate budgets, and task objectives:

1. On scaffold-, time-, and environment-held-out tasks, invariant-constrained candidates should have higher constraint-satisfaction rates and better-calibrated uncertainty than retrieval- or supervision-dominant AIDD. A null result or a baseline advantage would reject this prediction.
2. When target and molecule identifiers are randomized and retrieval is disabled, a language model restricted to parsing should lose less performance than a semantic-memory baseline. Equal or larger degradation would reject the claimed prior isolation.
3. At a fixed assay budget, an information-gain policy should reach a preregistered mechanism-discrimination criterion with fewer experiments or greater uncertainty reduction per cost than random search, standard Bayesian optimization, and LLM-only planning. No improvement would reject the efficiency claim.
4. In virtual-cell tasks with held-out energy budgets, membrane permeabilities, and perturbations, generated networks should satisfy a larger fraction of the applicable (*B, E, M, I, R, V*) predicates and recover more rapidly. Failure to generalize would reject the life-design hypothesis.

All four predictions require preregistered task packages, confidence intervals, and an explicit negative-result policy. They are not quantitative results of the present case studies.

### 8.5 Claim ladder for constructive evidence

Constructive outputs should carry an evidence level in the Living Lab Notebook: L0, formal specification; L1, simulator-consistent invariants; L2, independent implementation replication; L3, invitro assay; L4, in-vivo or organism-level validation; and L5, translational evidence. The current manuscript reports L0–L1 as a proposed protocol for this extension and reports only artifact-level validation for the implemented molecular and structural workflows. A higher-level claim is not permitted to inherit credibility from a lower-level simulation.

## 9 Discussion

BloClaw’s main contribution is the boundary between a language model and a scientific execution environment. Capability-state gating makes unsupported tasks visible before execution. Recovery events make operational failure inspectable. The Notebook turns a response into a structured record that can be reviewed and rerun. These properties address accountability and reproducibility, not only conversational fluency.

The constructive perspective changes the role of data without denying its value. In a conventional predictive workflow, historical data often define both the hypothesis space and the evidence used to score it. In the proposed mode, explicit invariants and mechanisms define the candidate space, while data are used for parameter calibration, model criticism, and reality checking. This separation makes it possible to ask whether a mechanism remains valid in an out-of-distribution environment instead of rewarding only similarity to the past. It also makes the source of novelty auditable: a result can be attributed to an axiom, a measurement, an executable model, or a human objective.

We use creator’s-eye design only as a metaphor for top-down specification of function and bottom-up composition under physical law. A first-principles mode cannot be literally prior-free: physical laws, chemical valence rules, simulators, model architectures, and even the definition of a desired phenotype are formalized priors. BloClaw’s proposed contribution is an epistemic contract that keeps generation, simulation, observation, and validation as distinct objects, so that no downstream claim exceeds the highest validated object.

The system should not be described as a complete AI drug-discovery platform. Complete ADMET prediction, de novo docking, protein folding, virtual-cell perturbation prediction, antibody affinity prediction, radiopharmaceutical modeling, peptide SAR, and autonomous robotic execution remain outside the validated scope declared here. They can be presented as migration targets or future work only after independent datasets, model versions, test sets, and biological validation are available.

## 10 Limitations, Ethics, and Reproducibility

The main limitations are the limited number of validated scientific capabilities, dependence on external tools and data quality, model and provider variability, and the absence of experimental or clinical validation in the present manuscript. There is no literal blank slate: laws, simulators, objectives, and formal representations are themselves prior knowledge, and the proposed constructive mode must quantify rather than hide those assumptions. The gap between first-principles constraints and organism-level biology also requires multiscale approximations and empirical calibration. The system must not be used to make clinical decisions or to present computational outputs as experimentally confirmed biological effects.

The current paper is a software/methods preprint with a complete system description, two regenerable case-study figures, a verified capability declaration, deterministic artifact checks, and an executable evaluation design. It does not report a fabricated benchmark, confidence interval, or biological discovery. For submission, the authors should archive the software release, environment specification, benchmark task package, prompts, logs, representative inputs, and output artifacts subject to licensing and privacy constraints.

## 11 Conclusion

This work presents BloClaw as a capability-gated AI4S workstation that connects natural-language task interpretation to controlled scientific execution, recoverable workflows, provenance recording, and human-verifiable deliverables. Its strongest current claim is architectural and operational: the system makes it explicit what can be executed, what evidence was produced, and what remains unvalidated. The proposed constraint-first constructive mode extends this foundation from retrospective prediction toward mechanism and viability-trajectory design: goal, invariants, mechanism, simulation, experiment, and evidence become one auditable loop. Quantitative claims about reliability, novelty, or reproducibility should be added only after the benchmark and ablation protocol in this manuscript has been run.

## A Reproducibility Checklist

- Software release and commit identifier: archive before submission.
- Model and provider configuration: record model ID, endpoint, generation settings, and date.
- Dependency environment: release a lockfile or container image.
- Benchmark task package: release inputs, expected outputs, and acceptance criteria.
- Prompts and task specifications: release the exact task definitions used in evaluation.
- Random seeds and repeated-run protocol: record all stochastic settings.
- Raw logs and output artifacts: archive task events, files, reports, and hashes.
- Human review procedure: predefine reviewer roles, criteria, and time measurement.

